# Emerging Activity Patterns and Synaptogenesis in Dissociated Hippocampal Cultures

**DOI:** 10.1101/2023.05.18.541345

**Authors:** Jyothsna Suresh, Mark Saddler, Vytas Bindokas, Anita Bhansali, Lorenzo Pesce, Janice Wang, Jeremy Marks, Wim van Drongelen

**Affiliations:** Department of Pediatrics, The University of Chicago, Chicago, IL 60637, USA; Committee on Computational Neuroscience, The University of Chicago, Chicago, IL 60637, USA; Department of Pharmacological and Physiological Sciences, The University of Chicago, Chicago, IL 60637, USA; Committee on Neurobiology, The University of Chicago, Chicago, IL 60637, USA

**Keywords:** **Net**work Maturation, Epileptiform Activity, Synaptic Detection Algorithm

## Abstract

Cultures of dissociated hippocampal neurons display a stereotypical development of network activity patterns within the first three weeks of maturation. During this process, network connections develop and the associated spiking patterns range from increasing levels of activity in the first two weeks to regular bursting activity in the third week of maturation. Characterization of network structure is important to examine the mechanisms underlying the emergent functional organization of neural circuits. To accomplish this, confocal microscopy techniques have been used and several automated synapse quantification algorithms based on (co)localization of synaptic structures have been proposed recently. However, these approaches suffer from the arbitrary nature of intensity thresholding and the lack of correction for random-chance colocalization. To address this problem, we developed and validated an automated synapse quantification algorithm that requires minimal operator intervention. Next, we applied our approach to quantify excitatory and inhibitory synaptogenesis using confocal images of dissociated hippocampal neuronal cultures captured at 5, 8, 14 and 20 days *in vitro*, the time period associated with the development of distinct neuronal activity patterns. As expected, we found that synaptic density increased with maturation, coinciding with increasing spiking activity in the network. Interestingly, the third week of the maturation exhibited a reduction in excitatory synaptic density suggestive of synaptic pruning that coincided with the emergence of regular bursting activity in the network.

## 1. Introduction

Dissociated hippocampal cell cultures display a stereotypical emergence of network activity patterns during the first weeks of development, ranging from low spiking activity at 5 days *in vitro* (DIV), increased spiking activity at 8 DIV, random bursting interspersed with spiking activity at 14 DIV and highly regular and synchronous network-wide bursting activity at 20 DIV that is reminiscent of an epileptic network (Wagenaar et al. 2006; Chiappalone et al. 2007; Pasquale et al. 2008; Charlesworth et al. 2015). Excitatory and inhibitory synaptogenesis, associated with the formation and reformation of axons, dendrites and synaptic contacts in the network play an important role in the network level manifestation of activity patterns in these cultures. Therefore quantification of the synaptic development may improve our understanding of the observed emerging neuronal network activity. To unequivocally quantify the density of synaptic connections in a neuronal network, one might apply electron microscopy (Van Huizen et al. 1985; Ichikawa et al. 1993; Papa et al. 1995; De Felipe et al. 1997; Boyer et al. 1998); however, quantification of synaptic density of an entire network at this extremely high-resolution is laborious and time consuming. Confocal microscopy techniques offer a promising method to capture and analyze high-resolution images of neuronal networks with marked synaptic structures (Durand et al. 1996; Zito et al. 1999; Mironova et al. 2007; Hohensee et al. 2008). To obtain quantitative information such as number of synapses from confocal images, the analysis is typically based on object-based colocalization of detected pre- and post-synaptic puncta structures. However, confocal images are prone to several noise components such as non-specific staining, auto-fluorescence of non-target structures and photon-noise which lead to spurious detections and random overlap between them when there is no real colocalization of synaptic structures. To mitigate the effects of these noise components, most existing synapse quantification strategies are predominantly manual or semi-manual (Glynn and McAllister 2006; Ippolito and Erogulu 2010; Danielson and Lee 2014) requiring user-set thresholds to distinguish signal from noise, which is extremely labor-intensive and introduces observer bias. Therefore tremendous amount of effort has been invested recently in developing automated approaches for detection of synaptic structures from confocal images. Most common automated procedures implement detectors that are based on arbitrary intensity threshold settings such as the mean and standard deviation of the image intensity levels (Schmitz et al. 2011; Schätzle et al. 2012; Harrill et al. 2015). While threshold-based image segmentation might eliminate low intensity background noise, the aforementioned noise components span wide range of intensities, resulting in different estimates of synapse counts for different threshold settings, leading to ambiguous results.

The goal of the present investigation is to quantify excitatory and inhibitory synaptogenesis at different functional stages of development in dissociated hippocampal cell cultures, ranging from sparsely connected networks that exhibit low levels of spiking activity to densely connected mature networks that exhibit periodic bursting. To quantify synaptogenesis, we evaluated and applied a novel automated, high-throughput approach, based on the spatial correlation between presynaptic and postsynaptic structures. This approach does not significantly depend on image intensity threshold and provides consistent estimation of noise components. We validated our approach using synthetic images and then applied it to quantify excitatory and inhibitory synaptogenesis during the first three weeks of maturation in dissociated hippocampal cell cultures. We found that there is an increase in excitatory synaptic density during the first weeks of maturation that coincided with increased spike activity at the network level. This was followed by a reduction in excitatory synaptic density towards the third week, suggesting that a synaptic pruning phase occurs as these cultures develop that coincides with regular bursting activity in the network.

## 2. Methods

We captured confocal images of neuronal cultures made from dissociated hippocampal neurons in rat embryos (E18). All experimental procedures involving animals were approved by and were in compliance with the Institutional Animal Care and Use Committee (IACUC) at The University of Chicago.

### 2.1 Cell cultures

Dissociated hippocampal neuronal cell cultures were prepared from embryonic day 18 Sprague Dawley rats as previously described (Suresh et al. 2016). Briefly, under deep ethrane anesthesia of the dam, E18 fetuses were extracted from the uterus and decapitated. The forebrains were removed, split sagitally in the midline and the meninges removed. Each hippocampus was gently freed from the surrounding cortex, minced and subjected to trypsin digestion followed by mechanical trituration to dissociate the cells. The dam was sacrificed under deep anesthesia by removal of the heart. The dissociated cells were seeded on poly-D-lysine coated glass cover-slips and multi-electrode arrays (MEAs) and maintained in neurobasal medium containing B27 supplement and GlutaMax (all from Life Technologies), in a humidified atmosphere (5% CO_2_, 95% atmospheric air at 37°C). Cytosine arabinoside (AraC) was added to the medium (2 µM at final concentration) at 4 days *in vitro* (DIV) to suppress the proliferation of non-neuronal cells such as glia. Two sets of cultures were used in the experiments, seeded at a density of ∼850 cells/mm^2^ and ∼650 cells/mm^2^ respectively. Cultures were maintained by replacing half of the media volume with freshly made culture media, every 4-5 days. The neurobasal medium contained (in mM): 51.7 NaCl, 26 NaHCO_3_, 0.91 NaH_2_PO_4_, 0.81 MgCl_2_, 5.33 KCl, 25 D-glucose, and 1.8 CaCl_2_.

### 2.2 Recording

#### Multichannel extracellular recording

Multichannel recordings were performed with multi-electrode arrays (MEA) and a MEA 2100 device (Multichannel Systems, Reutlingen, Germany). The MEAs contain 60 titanium nitride electrodes, laid out in a square grid: electrode diameter was 30 µm and inter-electrode distance was 200 µm. Experiments were performed in a controlled environment (5% CO_2_, 95% atmospheric air, and temperature 36 - 37°C). Each recording was done over a 15 min time period at a sample rate of 25 kHz/channel and a bandwidth of 1 Hz – 3 kHz. All recordings from the MEA were performed in neurobasal medium.

#### Intracellular recording

Standard electrophysiological recordings were obtained from the coverslips using whole-cell current-clamp technique under the visual guidance of a Axioskop 2 plus microscope (Carl Zeiss, Inc., Thornwood, NY, USA), connected to a Nikon CoolSnap HQ2 camera (Nikon Corporation, Tokyo, Japan) and imaged using Nikon Imaging Software (NIS Elements AR, Nikon Inc., USA). Patch electrode pipettes were fabricated from filamented borosilicate glass capillaries (Warner Instruments LLC, Hamden, CT, USA) using a P-97 micropipette puller (Sutter Instrument Company, Novato, CA, USA). The electrodes were filled with pipette solution containing (in mM): 140 K-gluconate, 10 HEPES, 2 MgCl_2_*6H_2_O, 4 Na_2_ATP, 1 CaCl_2_*6H_2_O and 10 EGTA (pH 7.3-7.4) with a resistance between 3 and 5 MΩ and all recordings were performed in extracellular artificial cerebrospinal fluid (ACSF) solution containing (in mM): 118 NaCl, 25 NaHCO_3_, 1 NaH_2_PO_4_, 1 MgCl_2_*6H_2_0, 3 KCl, 30 D-glucose, and 1.5 CaCl_2_. Neuronal activity was recorded using a MultiClamp 700B amplifier (Molecular Devices, Sunnyvale, CA, USA), and digitized and acquired at 25 kHz using a Digidata 1440A interface (Molecular Devices).

### 2.3 MEA Data Analysis

To quantify network activity we calculated the mean firing rate (spikes/sec) and mean burst rate (bursts/min) averaged across all electrodes in the MEA. Extracellular recordings were filtered off-line by a digital filter (a Butterworth filter, second order band pass 300 Hz - 1.5 kHz) using Matlab software (MathWorks, Natick, MA, USA). The filtered output was used to detect spikes, defined as negative deflections that exceeded five standard deviations of the filtered signal. The multi-unit spike trains were saved in rasters as arrays of 0’s (no spike) and 1’s. To detect bursts, the spike rasters were used as input to a leaky integrator with a time constant of 50 ms, a value close to the time constant of a hippocampal pyramidal cell (Staff et al. 2000). The output, which represents the integrated spike activity was used to detect bursts (van Drongelen et al. 2006; Suresh et al 2016). Burst detection threshold was set at four standard deviations of the integrated spike activity amplitude to identify the individual bursts.

### 2.4 Immunofluorescence

In order to quantify synaptogenesis, cultures were fixed using 4% paraformaldehyde in PBS and stained after 5, 8, 14 and 20 DIV. Excitatory and inhibitory synapses were stained in separate cover-slip preparations. To identify excitatory synapses, neurons were triple stained to label dendrites and excitatory pre- and post-synaptic terminals, using chicken-anti-MAP2 (Abcam 1.9µg/ml), rabbit anti-vGluT1 (Synaptic Systems, 10µg/ml), and mouse anti-PSD-95 (UC Davis/NIH NeuroMab Facility, 10µg/ml) respectively. To identify inhibitory synapses, neurons were triple stained to label dendrites and inhibitory pre and post-synaptic terminals, using chicken-anti MAP2 (Abcam 1.9 µg/ml), rabbit anti-vGAT (Synaptic Systems 10µg/ml), and mouse anti-gephyrin (Synaptic Systems 10µg /ml). Binding was detected with Alexa Fluor 488–labeled, highly cross-adsorbed goat anti–chicken IgY, Alexa-647-labelled highly cross-adsorbed goat anti–rabbit IgG, and Alexa-594-labelled highly cross-adsorbed goat anti-mouse IgG (1:500; Jackson). Cells were incubated in DAPI (300 nM for 2 min) to label nuclei, and mounted in Cytoseal-60 (Thermo Scientific).

### 2.5 Image capture

Cover-slips were imaged with a 63×, 1.40 numerical aperture, oil immersion objective on a laser scanning confocal microscope (Leica SP5 AOBS, in resonant scanner mode), with identical illumination acquisition settings across staining conditions. We used 12-bit dynamic range and highly-sensitive, linear HyD detectors for the pre- and post-synaptic puncta channels and standard illumination settings were created to make use of the dynamic range for the typical staining signals in this preparation. The image capture procedure is shown in Figure 1. For a given age of the culture (5, 8, 14 or 20 DIV) and each synapse type (i.e., excitatory or inhibitory), we fixed and stained cultures on two coverslips (Fig 1A). In each coverslip we sampled 81 locations to capture images, each image measuring 51.2µm × 51.2 µm (1024 × 1024 pixels) (Fig 1B). We thus captured a total of 162 images for each synapse type, for a given age within a culture. Sequential line scans were used to capture four separate channels of high resolution images of dendrites (Alexa 594), presynaptic puncta (Alexa 647), post synaptic puncta (Alexa 488) and nuclei (DAPI), using laser lines at 405 nm, 488 nm, 561 nm, and 633 nm, respectively. Figure 1C depicts a representative merged image (1024 × 1024 pixels) formed by superimposing the aforementioned four channels. Figures 1D and 1E depict zoomed-in sample images (200 × 200 pixels) capturing excitatory and inhibitory synapses respectively along with the dendrites.

**Fig 1.**
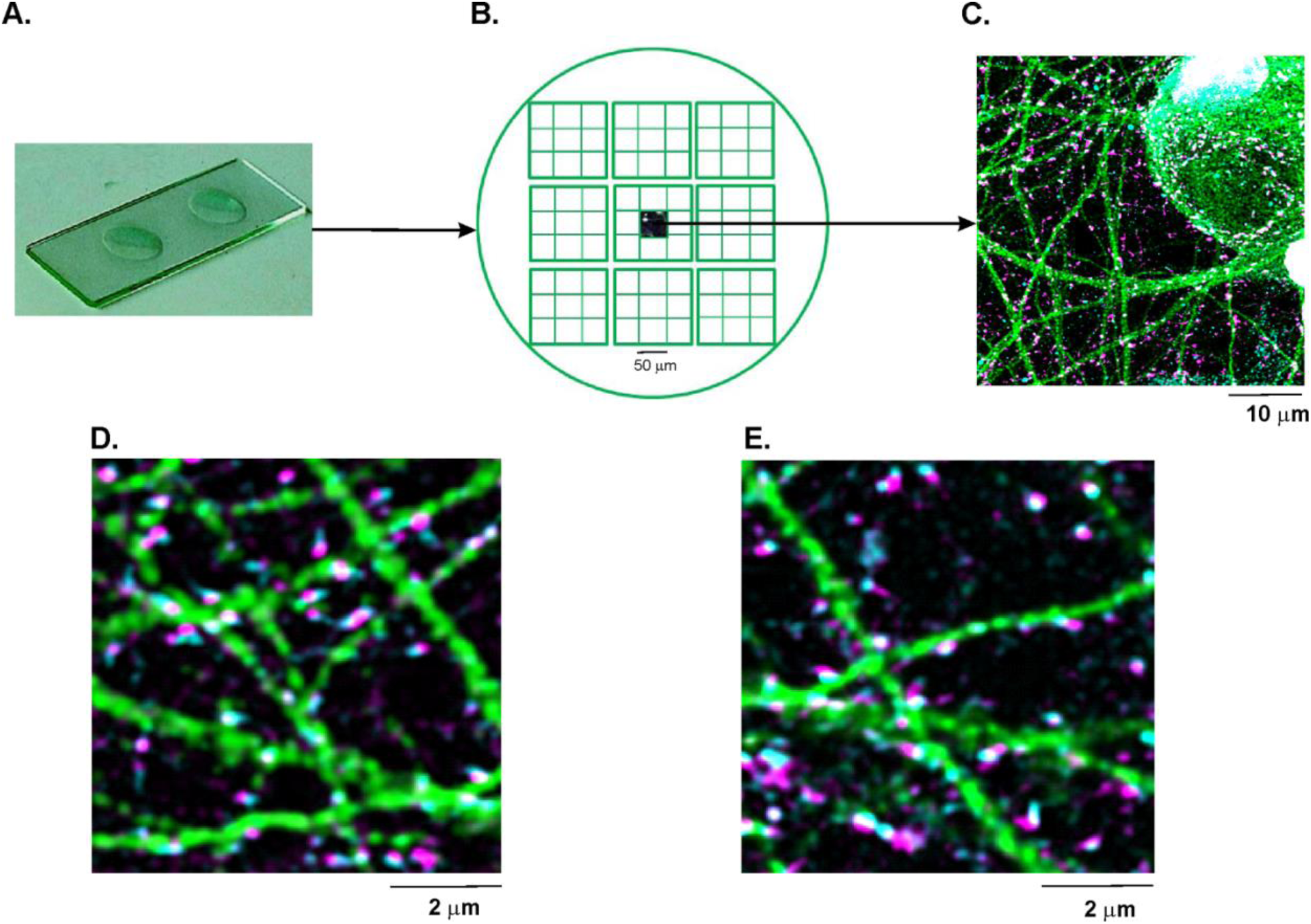
Immunocytofluorescence and image capture. A) Microscope slide showing two cover-slips that contain fixed and stained neuronal cell cultures. B) Cartoon of a single cover-slip, from which 81 multi-channel images were captured, arranged in a 9-by-9 grid. C) An example of one of the 81 multi-channel images, captured from the cover-slip. This is a merged image formed by superimposing four separate channels each capturing the pre-synaptic puncta (magenta), post-synaptic puncta (cyan), dendrites (green) and soma (grey). A zoomed in multi-channel image comprising of D) excitatory and E) inhibitory pre- and post-synaptic puncta as well as dendrites labeled with VGluT1/VGAT, PSD-95/Gephryn and MAP-2 respectively.

Digital emission filter windows were tuned to non-overlapping bands covering dye emission peaks. Raw multi-channel images were acquired in a Leica Image File (LIF) format, containing 12-bit grayscale data. Uniform grid capture was controlled by a template created in Matrix software (Leica), enabling autofocus and z-stack capture at each sample location. Since the dissociated hippocampal cell cultures used in the study are essentially two dimensional where the neuropil is <1µm high in these cultures, we collected 3 z-stacks, with a 0.33µm z-step size to cover that entire range. Thus, each image-stack contained images of three planes: one plane with the neuropil in focus, one focal plane above the neuropil and one focal plane below the neuropil.

### 2.6 Preprocessing and automated synapse detection

We performed an initial inspection prior to image analysis to discard images that were of poor quality where the signal was indistinguishable from background. We used a criterion of SNR < 0.5, as poor quality images. To remove distortion arising from the microscope’s point spread function and to improve signal-to-noise ratio, images were pre-processed using Huygens deconvolution software (Huygens v.4.5.1, SVI, Hilversum, The Netherlands) using maximum-likelihood estimation and signal to-noise ratio of 5. Subsequent image analysis involved automated synapse quantification of series of images, implemented in batch-processing mode. This was performed on a Windows 7 computer using a customized script written in ImageJ (https://imagej.nih.gov/ij/index.html).

Synapse identification was applied to the maximum intensity projections from the three z-planes, as our cell cultures are essentially a 2D monolayer of cells (neuropil thickness < 1 µm). We identified a synapse as colocalized pre- and post-synaptic puncta with their centroids lying within a distance of 250 nm, located on dendrites. This colocalization distance criteria of 250 nm was based on reports from super-resolution microscopy studies using rodent cortical cell cultures (Dani et al. 2010), that all relevant synaptic protein labeling can fit within a 250 nm radius from the midpoint of a synapse. Furthermore, electron microscopy studies have also shown that the dimensions of the pre- and post-synaptic puncta terminals are ∼ 200 nm-wide in diameter and separated by a ∼50 nm-wide synaptic cleft (Ribrault et al. 2011). The detection of synapses was performed in a three-step procedure described in the following.

<u>Step 1</u> involved the identification of putative pre- and post-synaptic puncta and dendrites from the raw images (Fig 2A). For the puncta channels (Fig 2A1-A2), we first applied a rolling-ball background subtraction to correct for unevenness in background illumination. We used a rolling-ball radius of 4 pixels, which would produce a protected zone of 9-pixel-wide diameter (450 nm) as sampled. We then used a 3×3 median filter, to remove point noise within the image. To enable detection of low intensity signals while removing the low intensity noise component, we employed a threshold at 45% of total intensity distribution which was below the mean intensity value commonly used (Schmitz et al. 2011; Schätzle et al. 2012; Harrill et al. 2015). We then captured the pixel locations of all intensity-maxima exceeding this threshold, and expanded each location by 2 pixels in all directions, creating 5-pixel wide (250 nm) puncta objects (Fig 2B1-2). Since we used local intensity-maxima to detect putative synapses, the threshold value of 45% of total intensity distribution proved best for our samples as it dropped several background local maxima thereby providing better signal to noise ratios. Next, we segmented the dendritic image by thresholding at mean pixel intensity to detect the dendrites (Fig 2B3), and dilated the dendritic mask by 2 pixels to capture all the colocalized pre- and post-synaptic puncta lying on and in close proximity to the dendrites.

**Fig 2.**
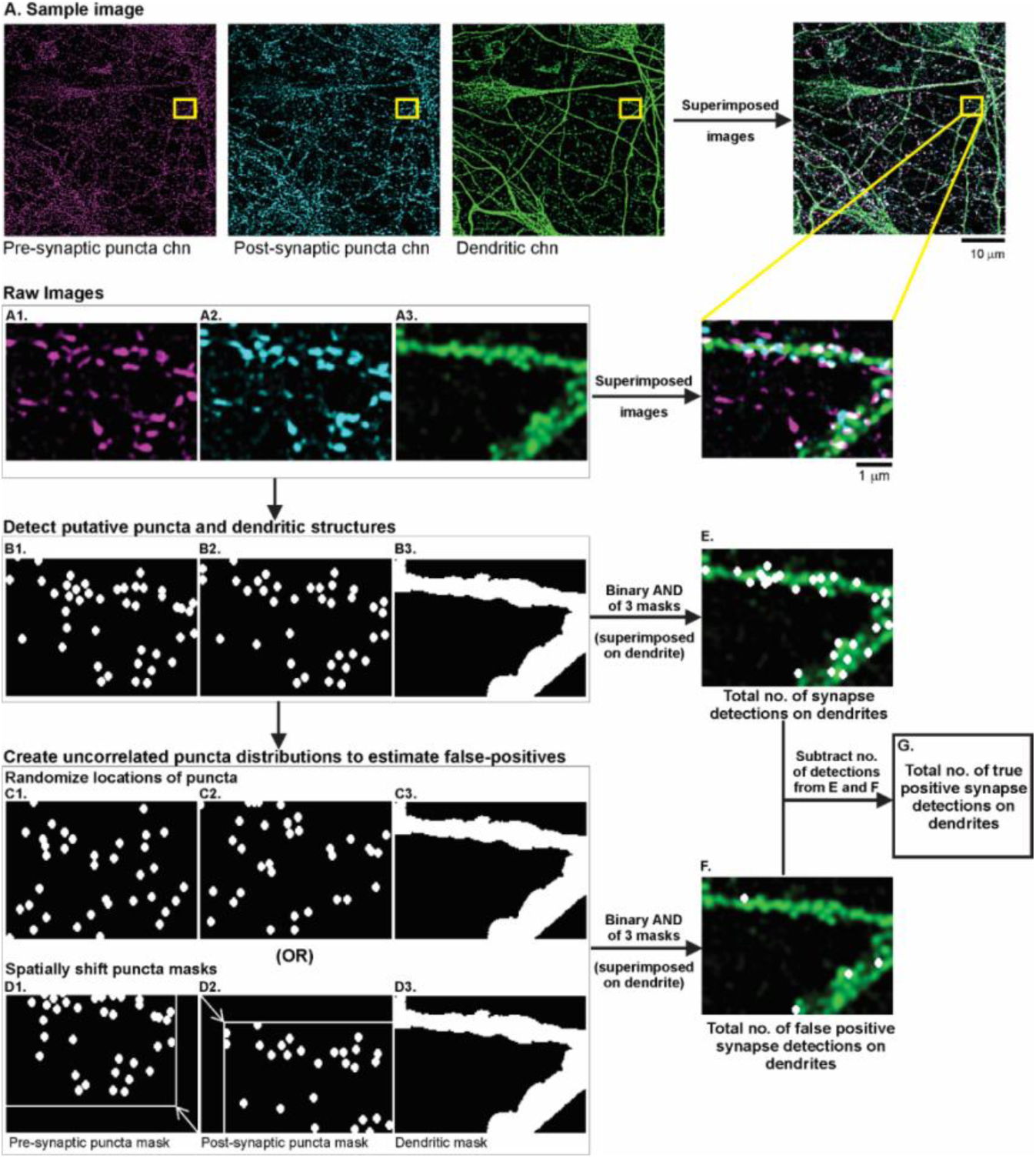
Flowchart of the automated synapse quantification algorithm. A) The first row depicts a sample raw image (1024 × 1024 pixels) labeled for the pre-synaptic structures (magenta), post-synaptic structures (cyan) and dendrites (green). The merged image formed by superimposing these separate channels, is shown in the rightmost panel in the first row. The yellow rectangle indicates the zoomed-in portion of the images used to explain the flowchart of our synapse quantification procedure. A1-A3) Raw images of the pre-synaptic puncta channel, post-synaptic puncta channel, dendritic channel respectively. B1-B2) Binary image of the pre- and post-synaptic puncta channels respectively, generated after implementing the following steps: rolling-ball background subtraction, 3×3 median filtering, thresholding at 45% of the total intensity distribution, identifying single-pixel local maxima, enlarging each maxima by 2 pixels in all directions, creating a 250 nm wide circular binary mask. Each circular mask represents a detected punctum. B3) Binary image of the dendritic channel, generated by setting a threshold at the mean pixel intensity in the raw image and extending the mask by 4 pixels. To estimate the noise due to random chance puncta-colocalizations on the dendrites, spatially uncorrelated puncta distributions were created using one of two independent methods: randomizing puncta locations in the binary masks (C1, C2) or spatially shifting the puncta masks relative to each other (D1, D2). E) Performing binary AND operation of the 3 original binary masks B1,B2, B3, gives an estimate of the total number of detections (colocalized puncta) on the dendrites. F) Performing binary AND operation of the 3 binary masks after creating spatially uncorrelated puncta distributions, gives an estimate of false positive detections on the dendrites. G) Subtracting the number of false positives (computed in F) from the total number of detections (computed in E) gives an estimate of the total number of putative synapses.

<u>Step 2</u> was to estimate the total number of colocalized pre- and post-synaptic puncta on the dendrites. To accomplish this, we performed a binary AND operation between the three binary masks generated for the dendritic, pre- and post-synaptic puncta channels (Fig 2B1-3) and counted the number (Fig 2E).

In <u>step 3</u>, we estimated the detection noise caused by random chance (or false positive) puncta-colocalizations on the dendrites. We repeated the AND operation of step2 after disrupting the spatial correlation between the pre- and postsynaptic puncta objects. This was achieved with two independent methods: (a) randomizing locations of the pre- and post-synaptic puncta within the respective images (Fig 2C1-2); (b) shifting the original pre- and post-synaptic masks relative to each other, i.e. spatial cross-correlation (Fig 2D1-2). To establish the reliability of our noise estimation procedure, the agreement between these noise estimates was assessed (see Fig 8). Finally, we computed the difference between the total number of detected colocalizations (in step 2, Fig 2E) and the noise estimate (Fig 2F) to get a noise-corrected estimate of the puncta-colocalizations (Fig 2G). Figure 3 depicts exemplary raw images (1024 × 1024 pixels) capturing the dendrites and the excitatory and inhibitory synapses as well as the corresponding binary masks.

**Fig 3.**
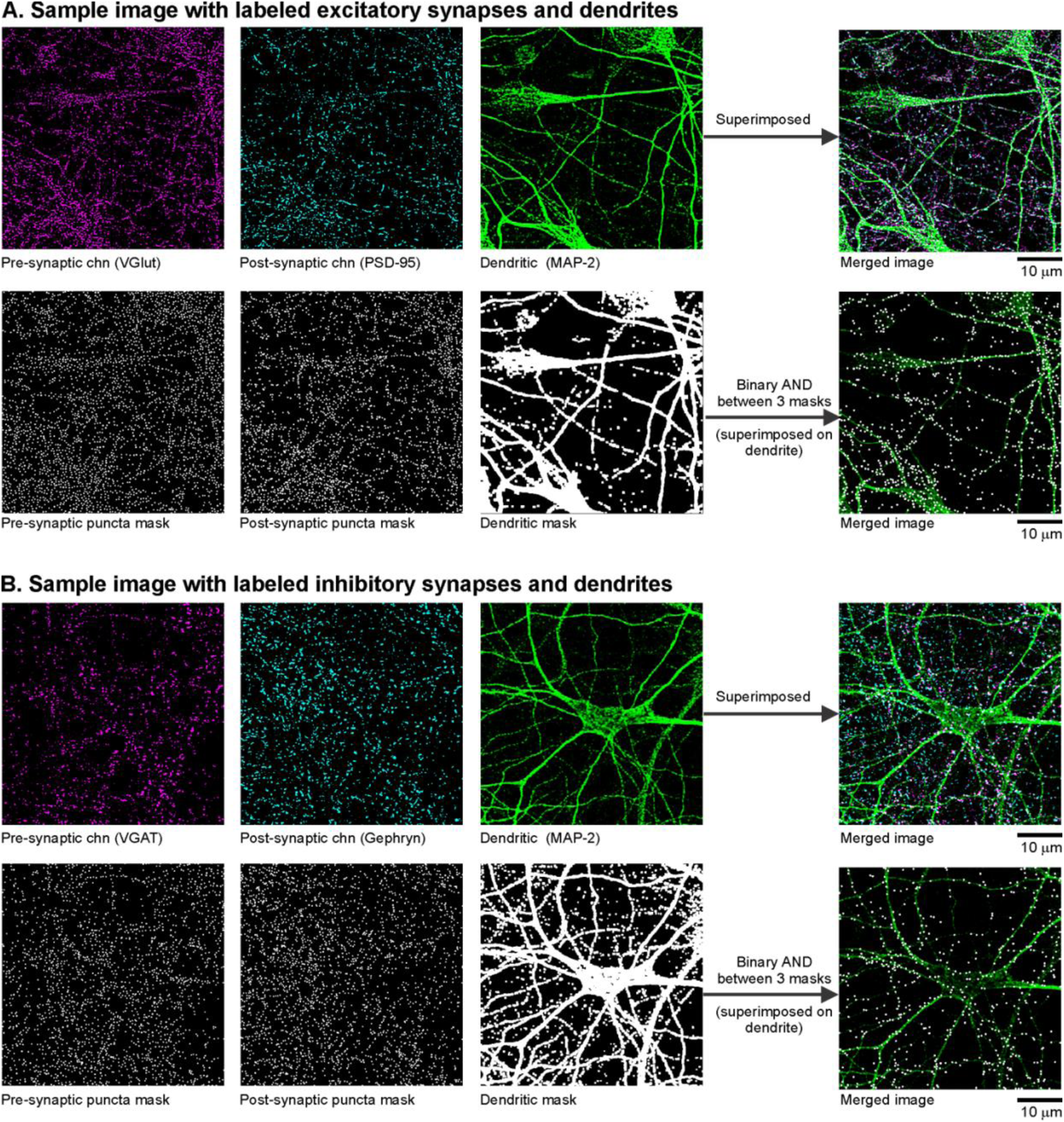
Binary masks generated from raw images. Sample raw images and the corresponding binary masks generated for the channels capturing the pre-synaptic terminals (magenta), post- synaptic terminals (cyan) and dendrites (green). Representative images are shown for A) excitatory and B) inhibitory synapses.

### 2.7 Validation of the synapse quantification algorithm using synthetic images

To evaluate the performance of our algorithm, we used it to detect signal and noise components in simulated binary images of dendrites, pre- and post-synaptic puncta channels (Fig 4). To mimic a situation of our experimental data, we created images 1024 × 1024 pixels in size, with 800 colocalized puncta on the dendrites (representing real synapses i.e. the signal) and added pre- and post-synaptic noise components. The objects in each puncta channel were depicted as 250 nm-wide circles (Fig 4A-B) and the dendrites were depicted as 5-pixel-wide lines (Fig 4C). The puncta objects representing real synapses (Fig 4D), were spatially correlated along the dendrites and were separated by a distance less than 250 nm. We created a series of images containing the original signal with different noise levels. This was done by adding different amounts of 250 nm wide circular objects, at spatially random locations in the simulated pre- and post-synaptic images. Using this procedure we obtained simulated image series with known signal and noise components, i.e. in contrast to the measured data, we had access to the gold standard. We then applied our algorithm to establish its performance using the known signal-to-noise ratio.

**Fig 4.**
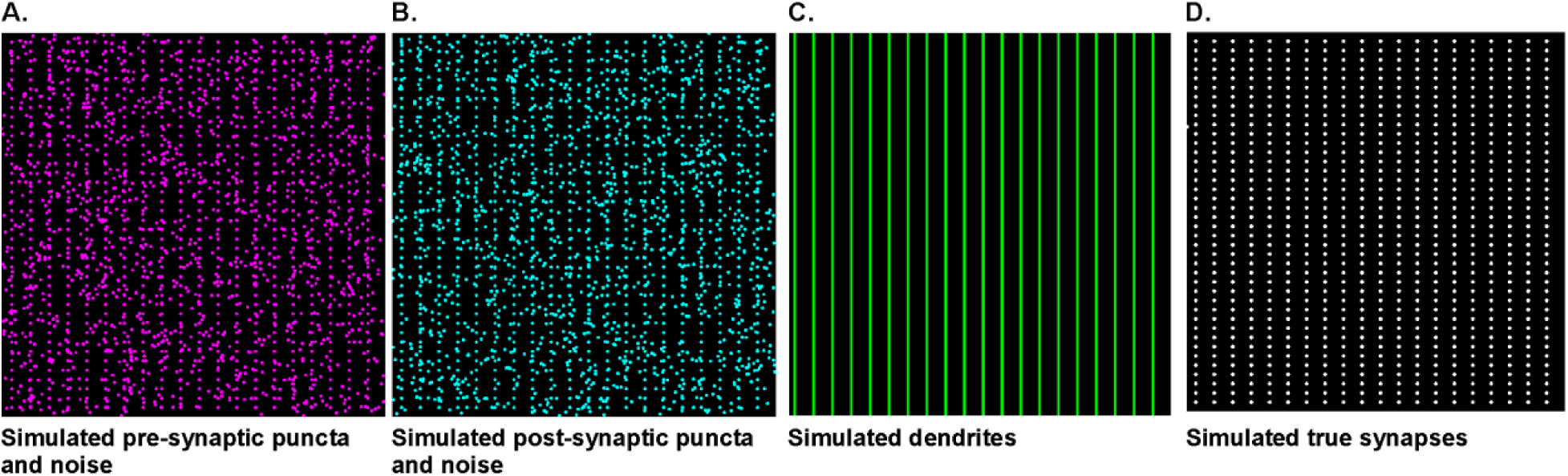
Simulated images used to validate synapse quantification algorithm. Simulated binary images of A) pre-synaptic puncta and B) post-synaptic puncta, each containing true synaptic terminals and a noise component. C) Simulated image of dendrites. D) Depiction of the true synapses (signal) simulated as colocalized pre- and post-synaptic puncta on the dendrites. These simulated images were used to test the performance of our synaptic quantification algorithm. Image dimension was the same as the experimental one, 1024 × 1024 pixels.

### 2.8 Statistical analysis

We quantified both excitatory and inhibitory synaptic densities across four developmental stages: 5, 8, 14, 20 days *in vitro* (DIV), in two independent series of cell cultures seeded at a density of ∼850 cells/mm^2^ and ∼650 cells/mm^2^ respectively. We analyzed a total of 2212 confocal images captured across all time points across both series of cultures (5 DIV, n=599; 8 DIV, n=548; 14 DIV, n=540; 20 DIV, n=525). The synaptic densities were calculated as the number of colocalized pre- and post-synaptic puncta per 100 µm^2^ dendritic area, after correcting for random chance colocalizations. We used the Data Analysis tool pack in Microsoft Excel 2013 (Microsoft, Redmond, Washington, USA), Matlab (MathWorks, Natick, MA, USA) and R (RDC Team, 2012) to compute basic statistics. Within each culture series, we used the nonparametric Wilcoxon rank-sum test to compare differences in excitatory synaptic densities across the developmental stages (5 vs 8, 8 vs 14 and 14 vs 20 DIV). The level of significance was determined at p<0.05 after corrections for multiple comparisons were applied using the Bonferroni method.

We analyzed the MEA cultures data using R (RDC Team, 2012). The longitudinal time dependent evolution of the neuronal activity over the DIV values was modeled using a mixed model for a nested design (electrodes within cultures) using the package lme4. Models of progressive complexity were tested using the package ANOVA test until adding terms produced no significant improvement in the model. We started with two random factors (electrodes nested within cultures) and a mean value and ended with a cubic fit (the cubic fit was adopted because the plots showed a clear increase in slope before decreasing again, which requires a third order term to be modeled). We kept only slope as a random factor in the fit because the biology of the models requires a zero intercept and adding higher order terms makes the model too flexible and hard to fit and interpret. Significance was set at 0.01 to account for testing of multiple quantities (4). To confirm the results, a nested bootstrap approach (random sampling of cultures with replacement, followed by random sampling of electrodes with replacement within those cultures – the electrodes were selected once for all the DIV values to preserve the longitudinal sampling of the experiment). Bootstrap results are reported as 99% confidence intervals (also to account for multiple comparisons).

## 3. Results

In this section we first present the typical network activity patterns that spontaneously emerge during the first three weeks of culture maturation. Next we quantify synaptogenesis during this period and validate our novel synapse detection algorithm. We especially determine robustness of our detection procedure with respect to the presence of noise and detector threshold selection.

### 3.1 Activity Patterns during Network Growth

Typical samples of the activity patterns recorded from dissociated cell cultures grown on MEAs, during the first three weeks of development are depicted in Figure 5. Although culturing the cells in neurobasal medium and in the absence of glia renders them relatively spine-free (Lesuisse and Martin 2002; Meberg and Miller 2003), our electrophysiological recordings have revealed vigorous activity associated with synaptogenesis (Fig 5). The extracellularly recorded spike activity (Fig 5A) is initially low or absent (5 DIV) and develops into low to moderate levels of spiking around the end of the first week of maturation (8 DIV). Towards the end of the second week (14 DIV), the network activity is dominated by irregular population bursts interspersed with spiking activity. This pattern of burst activity finally transitions into an extremely regular bursting pattern with very few or no extra-burst spikes (20 DIV). We quantified this activity in terms of mean spiking and bursting rates (Table 1) and found that there was measurable spiking and bursting activity by 8DIV which increased significantly by 14 DIV, followed by a reduction towards 20 DIV. The corresponding intracellular measurements confirm the activity patterns observed in the MEA (Fig 5B). The bottom traces in Figure 5B show the details of the measurement marked by the horizontal lines in the top traces of this panel. The sample of the initial state at 5 DIV is associated with subthreshold fluctuations. Action potentials are observed a few days later (8 DIV) and they start to cluster into occasional bursts at 14 DIV. Interestingly, the regular bursting we observe at 20 DIV is clearly associated with paroxysmal depolarization shifts (PDSs), a cellular hallmark of epileptiform activity (Fig 5B). The structural development as observed in the confocal images is depicted in Figure 5C. Although individual synaptic structures are not included in these images, the growth of the neurites involved in network maturation can be observed. At 5 DIV, the areas covered by the neurites of the individual neurons hardly overlap, while the density of the neurites clearly increases with time in the images at 8, 14, and 20 DIV. These observed functional and structural developments during the first three weeks of maturation motivated us to quantify synaptogenesis in during this epoch.

**Fig 5.**
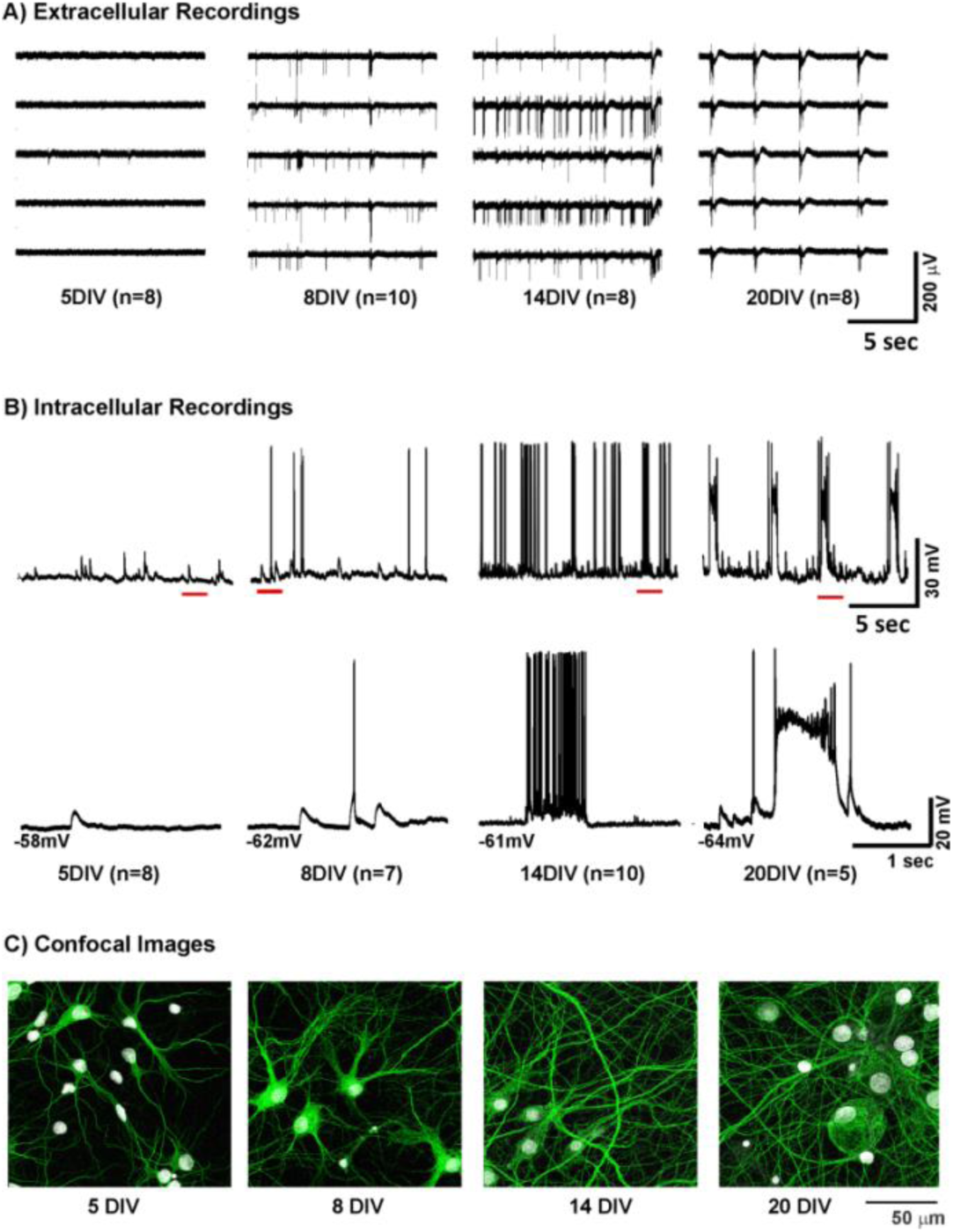
Activity patterns during network growth. A) Representative samples of network activity observed across the MEA at 5, 8, 14 and 20 DIV. This culture was seeded at a density of 650 cells/mm^2^. The extracellular data shows initial low levels of activity (5 DIV), evolving into irregular spiking pattern (8 DIV). Occasional bursts are observed at 14 DIV, while regular bursting emerges at 20 DIV. B) Intracellular recordings across the same stages of network maturation show that the initial activity at 5 DIV typically consists of subthreshold fluctuations while occasional spikes are observed a few days later (8DIV). Interestingly, the bursting that emerges after 14 DIV consists of typical grouped spikes. However, the regular bursting pattern around 20 DIV is characterized by paroxysmal depolarizations, a cellular hallmark of epileptiform activity. C) Development of hippocampal networks *in vitro*. Images of dissociated hippocampal neuronal cultures fixed and stained on coverslips at 5, 8, 14 and 20 DIV, at a density of 650 cells/mm^2^. Each image in this depiction is a mosaic formed by stitching 9 individual sample images (each 51.2µm×51.2µm), laid in a 3-by-3 grid (Fig 1). Merged images shown here comprise of dendrites and cell bodies (labeled in green and grey colors respectively).

**Table 1.** Tracking network activity during development in hippocampal cell cultures. The developmental stages tracked in this study are 5, 8, 14 and 20 days *in vitro* as indicated in the first column. The number of experiments (*n*) corresponding to each developmental period is indicated in the second column. The respective spike rates (spikes/sec) and the burst rates (in bursts/min) are expressed as mean ± SEM.

|  | Days *in vitro* | (*n*) | Spike Rate (spikes/sec) | Burst Rate (bursts/min) |
| --- | --- | --- | --- | --- |
| 716 |  |  |  |  |
| 717 |  |  |  |  |
| 718 | 5 | 8 | 0.38±0.01 | 0.25±0.06 |
| 719 | 8 | 10 | 1.65±0.07 | 4.13±0.06 |
| 720 | 14 | 8 | 4.83±1.25 | 23.50±0.87 |
| 721 | 20 | 8 | 3.93±0.12 | 16.25±0.78 |

### 3.2 Quantification of Synaptogenesis

Because we developed a novel algorithm to automate quantification of synaptic structures, we started with evaluating its performance on simulated images. Next, we used the approach to quantify synaptic structures from confocal images of hippocampal cultures captured at 5, 8, 14 and 20 DIV.

#### Simulated Images

We produced simulated images with known signal-to-noise ratios (SNR) (Methods, Fig 4) to determine the performance of our synapse detection algorithm under realistic, noisy conditions. Figure 6 depicts the relationship between the real number of synapses, noise levels and their estimates. We found that our detector reports a consistent estimate of number of puncta-colocalizations (green solid line, Fig 6) for SNR values between 1.5 and 6, which is the relevant range for our measured data set. Thus, in this range, the sensitivity of our detector is rather stable, i.e. 90-92% (the ratio between the detected and true synaptic structures shown by the green lines in Fig 6). In contrast, the error in the synaptic density estimation without noise correction, due to false positive detections (Type I error), increases significantly (10-40%) with decreasing SNR values in this range (purple line in Fig 6).

**Fig 6.**
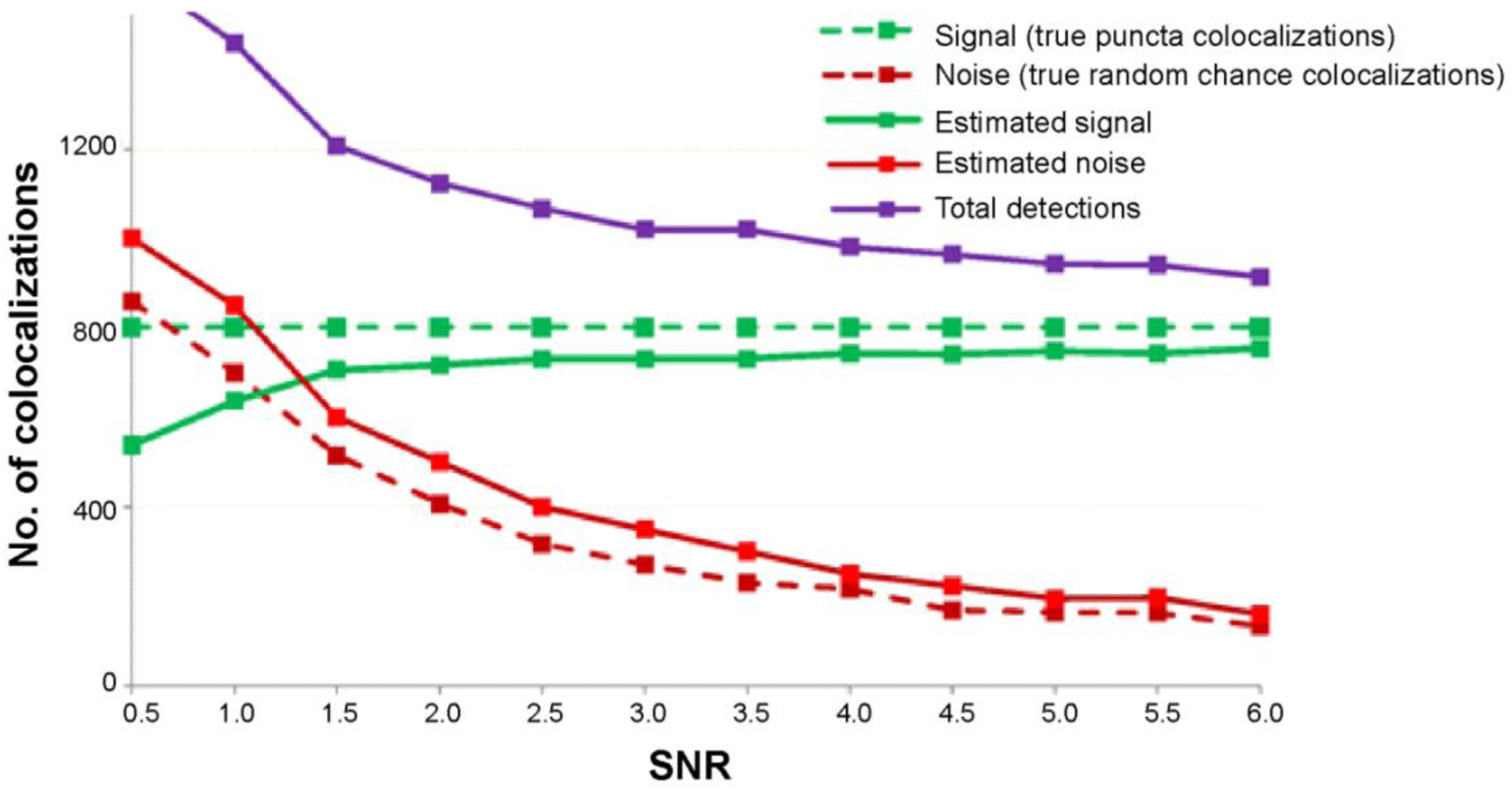
Performance of synapse quantification algorithm using simulated images. Detection as a function of signal-to-noise ratio (SNR). In the simulated images, signal is the number of real puncta-colocalizations (800 in this example) and noise is the number of random chance colocalizations. It can be seen that with decreasing SNR (i.e. increase in noise level), both the total number of detections (purple line) and noise estimate (red solid line) both increase, effectively keeping the number of true positive detections (green solid line) almost constant. For SNR values between 1.5 and 6, the algorithm performs with 90-92% sensitivity in estimating the number of colocalizations, after which it starts to severely underestimate these numbers. The true signal and noise components in the simulated images are shown by the dashed green and red lines respectively.

#### Hippocampal Cultures

Using the confocal images, we quantified excitatory and inhibitory synaptogenesis in two series of cultures that were plated at different cell densities: 650 cells/mm^2^ and 850 cells/mm^2^ (Fig 7A). In both series, we found that excitatory synaptic density initially increased and reached a peak around 8-14 DIV and then decreased towards 20 DIV (Fig 7B-C). Since there was no clear indication of neuron apoptosis (Fig 7A) during the later developmental stages, this observation of overshoot followed by a reduction in excitatory synaptic density is suggestive of synaptic pruning. On the other hand, density of inhibitory synapses showed an increasing trend as the cultures matured. Interestingly, the culture that was plated at sparser cell density showed onset of synaptic pruning at a later stage in the development as compared to the culture plated at a higher cell density (Fig 7B-C).

**Fig 7.**
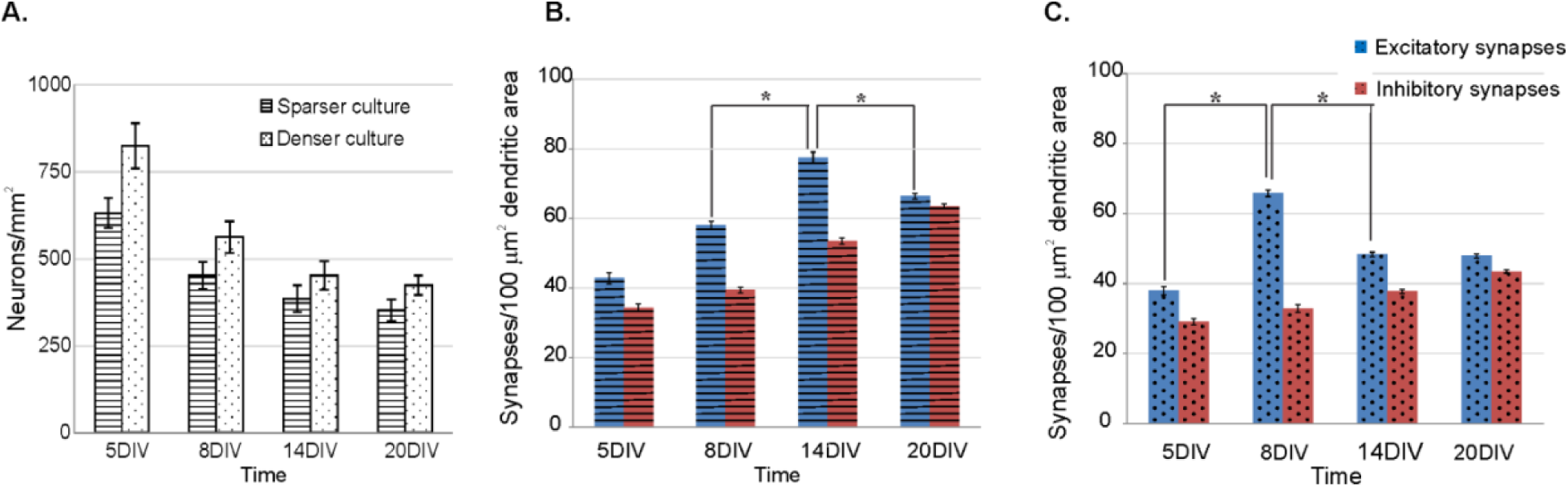
Excitatory and inhibitory synaptogenesis in hippocampal networks. (A) Comparison of cell densities in two series of dissociated hippocampal cell cultures used in the study: a sparser culture plated at 650 neurons/mm^2^ and a denser one plated at 850 neurons/mm^2^. The two panels on the right show the excitatory and inhibitory synaptic densities of the sparser (B) and denser (C) culture across developmental stages. Synaptic densities are presented in terms of mean ± SEM. Note that as the culture matures, excitatory synaptic density increases, exhibits a maximum followed by a decrease in density. Inhibitory synaptic density on the other hand exhibits an increasing trend. Furthermore, the maximum of the excitatory synaptic density is observed later for the sparser culture (14 DIV) as compared to the denser one (8 DIV). Statistics were computed using Wilcoxon rank-sum test. * Indicates statistical significant difference below an adjusted 0.05 level after Bonferroni correction.

#### Estimation of Detector Noise

In contrast to the simulated images, signal and noise levels arising from random colocalization are unknown in the experimental data. To compensate for this lack of knowledge (in part), we estimated and compared the noise levels in the detector output with two independent methods: randomizing the locations of the binary puncta-objects in the masks, and spatially shifting the original masks, i.e. spatial cross-correlation analysis. We found that these independent estimates were in agreement. A representative example of our noise estimates is shown in Figure 8. Figure 8A depicts the results obtained from a series of image samples in a single coverslip, demonstrating that the detector behaved consistently across the samples in a single coverslip. Figure 8B plots the spatial cross-correlation function that asymptotes to the estimated noise level for one of the samples. Noise estimates obtained with both methods were similar, a typical result for image#81, estimating a noise level of 25.4 synapses/100 µm^2^ dendritic area, is shown in Figure 8 (dashed line).

**Fig 8.**
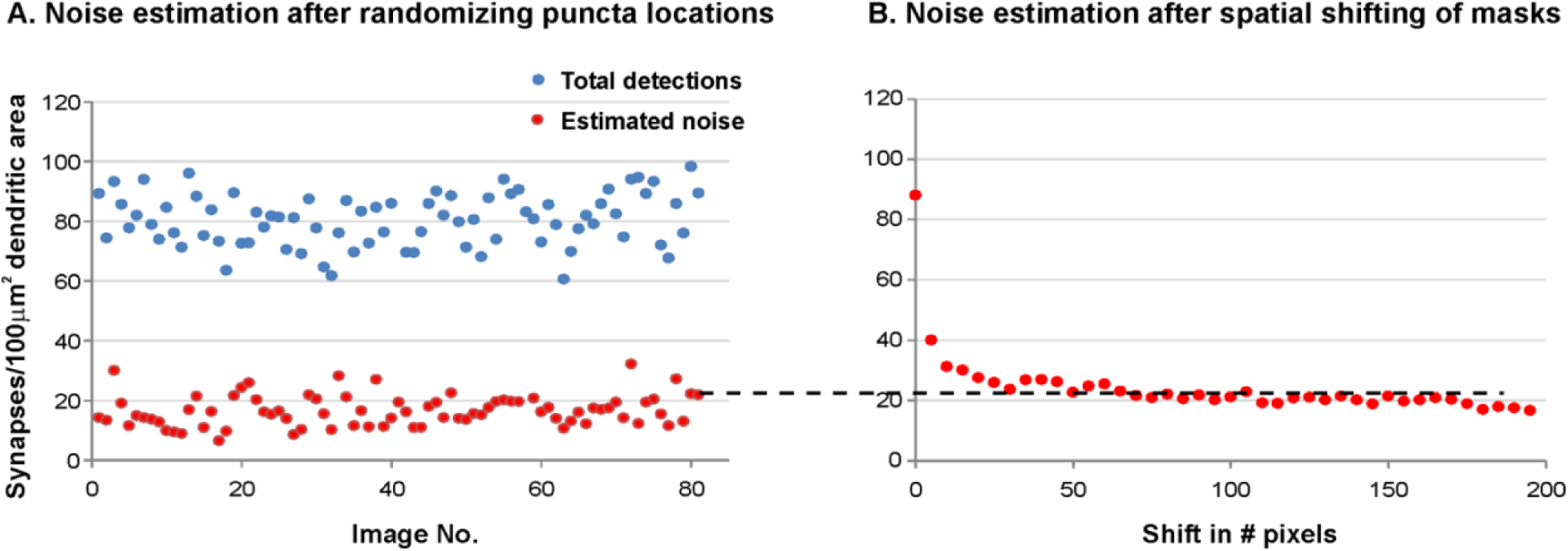
Noise estimates obtained from randomizing or spatial shifting of binary puncta. (A) Results obtained from randomizing the locations of the binary puncta. Blue and red dots represent the estimate of total number of puncta colocalizations and the estimate of noise, due to random chance puncta colocalizations. Each point on the x-axis represents an individual image sample collected from a single cover slip. (B) Result from the spatial cross-correlation function. Red dots represent the puncta colocalizations, with values asymptoting to the noise estimate (dashed line). Note that the noise estimates of both procedures, here shown for image#81, are similar.

#### Robustness of the Detector

One important component of an automated quantification of synaptogenesis is the procedure to set the detection threshold for potential synaptic elements. Usually, detector output critically depends on the threshold level. The salient aspect of our novel synapse quantification algorithm is that irrespective of the threshold set on fluorescence intensity of the pre- and post-synaptic puncta channels, the final synaptic density calculated is rather consistent. This property is demonstrated in Figure 9, where each of the four sub-plots depicts synaptic densities computed from representative images captured at 5, 8, 14 and 20 DIV. In each plot, the detected dendritic synaptic density (corrected for random chance puncta colocalization) is plotted versus the threshold settings used on the pre- and post-synaptic puncta channels. It can be seen that the estimate of synaptic density is similar for intensity thresholds up to 65-70% of the overall intensity distribution. Unsurprisingly, for higher threshold settings, we observed a drop in the detected synaptic density due to failure of detection of low intensity signal.

**Fig 9.**
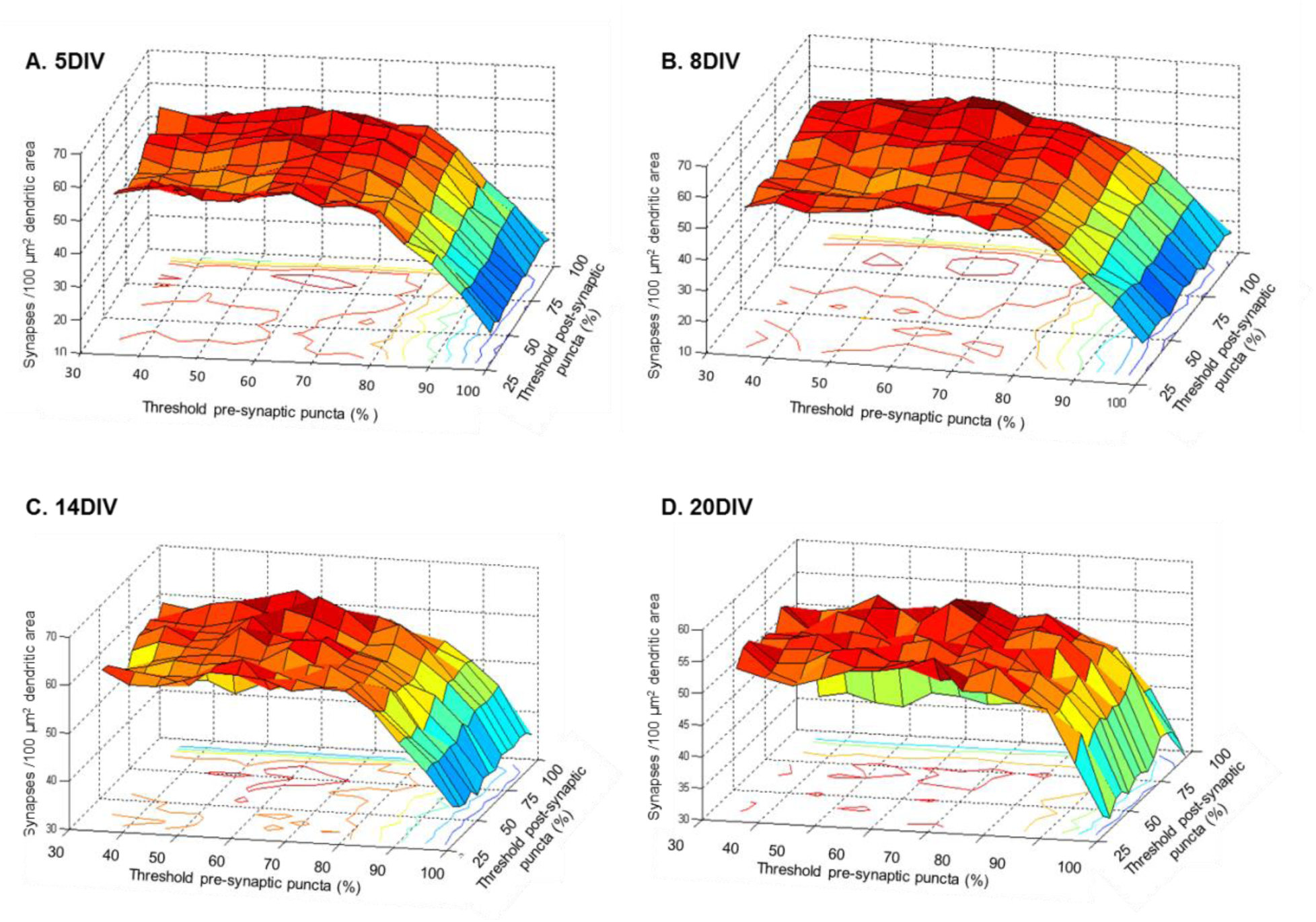
Effect of threshold on synapse quantification. Each panel (A-5 DIV, B-8 DIV, C-14 DIV, D-20 DIV) depicts the effect of threshold forW the pre- and post-synaptic puncta detection (horizontal axes) on the estimate of synaptic density (vertical axis). The threshold intensity values are set as the percentage of overall pixel intensity distribution. Synaptic density estimates were corrected for random chance puncta colocalization.

## 4. Discussion

In this work, we applied and evaluated a novel method for automated detection and quantification of synapses in confocal images of neuronal networks and applied it to quantify excitatory and inhibitory synaptogenesis in dissociated hippocampal cell cultures. Using both simulated and experimental data sets, we established that the automated procedure provided consistent estimates of synaptic density, and that these estimates were rather independent of noise contamination of the images and detector threshold selection (Figs 6, 8 and 9). In the experimental data, we found that excitatory synaptic density increased and reached a peak around 8-14 DIV, and then declined towards 20 DIV, which might be suggestive of synaptic pruning (Fig 7B,C). On the other hand, we found that the density of inhibitory synapses increased as the culture matured. Our observations of excitatory and inhibitory synaptic development could be simulated with a simplified activity-dependent network growth model (Fig 11).

Our motivation behind this study was to correlate the emergence of structure and function in developing neuronal networks in vitro. Figs 5A and 7B depict the emergence of functional activity and structural connectivity respectively at specific developmental stages of hippocampal cell cultures (seeded at a density of 650 cells/mm^2^). Based on these observations, we can speculate how synaptogenesis in these cultures might shape network activity during maturation. It can be seen that by the beginning of the second developmental week in vitro (8DIV), the initially isolated neurons develop axons and dendrites to form random network connections via synapses as reflected by the gradual increase in excitatory and inhibitory synaptic densities. This is manifested at the functional level in the form of initially low spiking activity (5 DIV) which then develops into measurable levels of spiking and bursting around the end of the first week of maturation (8 DIV). By the end of second week in vitro (14 DIV), although there is a continued increase in inhibitory synaptic density, there is a significantly large increase in excitatory synaptic density. This could lead to a massive excitatory drive to a large number of target neurons that might not be sufficiently contained by the inhibitory connections. This is manifested at the network activity level in the form of high levels of bursting and spiking activity (Table 1). Towards 20 DIV, however, there is a reduction in excitatory synaptic density along with a parallel increase in inhibitory synaptic density that coincides with a reduction in mean spike and burst rates (Table 1) at the network level. The spontaneous activity at this stage is characterized by regular alternating periods of bursting and quiescence (Fig 5A).

Although many groups have investigated synaptogenesis in *in vitro* cell cultures, a direct comparison of our results with theirs is difficult because of different experimental procedures (Catherine Croft Swanwick 2006 confirms inhibitory synaptogenesis). For example, we used glia-free and serum-free cultures maintained in neurobasal media with B27 supplement, which render them relatively spine-free (Lesuisse and Martin 2002; Meberg and Miller 2003). Other groups have used neuron-glia cultures maintained in serum-based medium, which develop dendritic spines [Van Huizen et al. 1985; Ichikawa et al. 1993; Schätzle et al. 2012; Harrill et al. 2015]. Furthermore, there are differences in seeded cell densities. In addition, different techniques, i.e. electron microscopy and confocal microscopy were applied to assess synaptogenesis. In spite of the differences in culture preparation and measurement technique, our data suggesting synaptic pruning is in agreement with the electron microscopy results reported by Van Huizen et al. (1985) and Ichikawa et al. (1993). Interestingly, both these studies used similar cell density as ours (∼900 cells/mm^2^). In contrast, the confocal microscopy results from Schätzle et al. (2012) and Harrill et al. (2015) did not find clear evidence of a pruning phase but rather an increase in densities during the first three weeks of maturation. Schatzle et al. used neuronal cultures in the presence of glia, while Harill et al. used a glia-free culture just as we did. However, as compared to our cell densities (650 cells/mm^2^ and 850 cells/mm^2^), both these studies employed a much lower cell density: 40 cells/mm^2^ and 315 cells/mm^2^ respectively. In conclusion, the applied culture density is the common difference between the studies that report pruning and those that do not. These findings are not necessarily contradictory since our experimental and modeling results demonstrate that the timing of the pruning phase in the synaptic development critically depends on the network’s cell density (Fig 7). We find that a low cell density is associated with a delayed pruning phase. Thus in the studies that employed low cell densities, e.g. Schätzle et al. (2012) and Harrill et al. (2015), the pruning phase could have occurred outside the window over which the culture was observed. We would also like to point out that we only observed and reported the synaptic densities at four discrete time points within a 3-week developmental period. Therefore, any significant changes in development of synaptic densities that could have occurred in between these time points were not captured in this study.

One problem of using confocal studies to quantify synaptogenesis is the fact that the staining procedures are not 100% specific for the target synaptic structures. For the synapse detection this results in spurious staining, leading to a significant noise component in the images. At the data acquisition level, uncertainty caused by this noise component can be reduced by staining both pre- and post-synaptic terminals, and by using colocalization of these structures as the synapse detection criterion. However, since both the pre- and post-synaptic terminal staining include spurious results, their combination will still create noise in the form of random chance colocalizations. Another problem with the identification of synaptic structures in the images is the presence of auto-fluorescence of non-target structures and therefore some degree of false detection associated with the intensity threshold of the detector. Different authors have developed strategies to estimate the noise components arising from random chance colocalization (Van Steensel et al. 1996; Lachmanovich et al. 2003; Costes et al. 2004). Similarly, several authors have attempted to circumvent the detection of non-target synaptic structures by using manual inspection of the images (Glynn and McAllister 2006; Ippolito and Eroglu 2010) or automated detection procedures that use an arbitrary intensity threshold value based on the mean and standard deviation of the image intensity level (Schätzle et al. 2012; Harrill et al. 2015). As with any detection process, the level of an arbitrary intensity threshold is a trade-off between missing identification of real structures (Type II error), and erroneous detection of noise components (Type I error). The goal of our automated synapse quantification approach was to mitigate, as much as possible, the problems created by the noise components and threshold effects (Figs 6, 8 and 9). We accomplished this by obtaining reliable estimates for the noise component in our images (Fig 8), resulting in a highly sensitive detector (90-92%) over a large range of SNRs (Fig 6). This reliable noise estimate made threshold selection a less critical property (Fig 9), which enabled us to employ a lower intensity threshold value (i.e. below the mean intensity) as compared with the mean pixel intensity that is commonly employed in existing detection procedures (Schmitz et al. 2011; Schätzle et al. 2012; Harrill et al. 2015). Our approach to synapse quantification is to use the spatial correlation of pre- and post-synaptic puncta along a neuronal surface i.e., dendrites in our case. Our current method of performing the AND operation on both the puncta and the surface masks would equally apply for other staining strategies in other types of cultures. For example, our approach could be extended to analyze images where the masks capture dendritic spines as well.

One finding of particular interest is the presence of a pruning phase during the development of the network (Fig 7). Albeit at a different timescale, several human studies (Huttenlocher 1979; Huttenlocher 1986) as well as other primate studies (Bourgeois and Rakic 1993; Wolff et al. 1995; Mimura et al. 2003), have also shown evidence for initial overproduction followed by pruning of neurites and synaptic structures. This similarity between the experimental data and reported clinical findings, as well as simulation studies, indicates that the dissociated cortical culture may be a useful model to study the rules underpinning synaptogenesis. Therefore, these *in vitro* models may ultimately help to understand synaptic mechanisms governing the development of the connectivity in neuronal networks.

## Grants

This work was supported by National Institute of Neurological Disorders and Stroke Grants R01-NS-095368 and R01-NS-084142.

